# Poly(A) probe HCR RNA-FISH specifically marks pyriform nurse cells in the brown anole lizard ovary

**DOI:** 10.1101/2025.09.10.675451

**Authors:** Zoe B. Griffin, Bonnie K. Kircher, Richard R. Behringer

## Abstract

Hybridization chain reaction RNA-fluorescent in situ hybridization (HCR RNA-FISH) is a powerful and increasingly used method for visualizing gene expression in cells and tissues. A probe set against polyadenylated RNA (poly(A)) is often used as a positive control for RNA integrity and staining quality. While optimizing this technique in the ovary of the brown anole lizard (*Anolis sagrei)*, we found that the poly(A) probe produced a strikingly specific and intense signal in pyriform cells, a specialized lizard-specific nurse cell type. This staining pattern was found in both whole-mount samples and paraffin sections, suggesting that poly(A) signal intensity can serve as a robust molecular marker for this cell type. The specific and robust signal facilitated segmentation of volumetric data to create the first 3D models of pyriform cells and quantify their sphericity. We also observed unusually diffuse DAPI staining in pyriform cell nuclei, distinguishing them from surrounding granulosa cells, pointing to possible differences in chromatin structure or nuclear organization. Together, these findings highlight the potential of poly(A) probes used in HCR RNA-FISH not only as a technical control, but also as a tool to selectively label specific cell types with high transcriptional activity or storage of abundant poly(A) transcripts.

## Introduction

The ovary is a complex and dynamic organ central to female reproductive biology. The primary role of the ovary is the maturation of oocyte-containing follicles for ovulation and the production of hormones for reproductive organ function. Oogenesis involves the progressive development of follicles, containing oocytes surrounded by somatic cells, which support the growing oocyte through signaling, structural support, and nutrient exchange in a complex and heterogeneous cellular environment (Barresi and Gilbert, 2020).

In reptiles, reproductive strategies are highly diverse, including oviparity (egg laying), viviparity (live birth), and ovoviviparity (development in eggs within the mother followed by live birth). There are also variations in traits such as clutch size, egg laying frequency, and reproductive modality, offering key insights into vertebrate evolution (Kunz and Orrell, 2004). Despite this diversity, the anatomy of the reproductive tract is well conserved across reptiles (Blackburn, 1998; Girling, 2002).

The brown anole (*Anolis sagrei)*, a small egg-laying lizard native to the Caribbean and now a widespread invasive species in the southeastern United States, has become an increasingly important model for development and reproductive studies due to its short reproductive cycle, well-defined embryology, ease of laboratory maintenance, and growing availability of genetic and imaging tools (Sanger et al., 2008; Sanger and Kircher, 2017; Rasys et al., 2019; Geneva et al., 2022; Kircher et al., 2024a). Female anoles lay a single egg at a time, approximately once per week, and alternate ovulation between the left and right ovaries (Jones et al., 1983; Kircher et al., 2024b). Females have two fully independent reproductive tracts, each consisting of an ovary, infundibulum, glandular uterus, and nonglandular uterus, with both sides converging with the digestive tract at a single cloaca (Kircher et al., 2024b).

The anole ovary is composed of numerous growing follicles of progressive size, from small previtellogenic follicles to large vitellogenic follicles with associated yolk (Ortiz and Morales, 1974; Kircher et al., 2024b). In squamate reptiles (lizards and snakes), each follicle contains a central oocyte surrounded by somatic cells (García-Valdez et al., 2019). These somatic cells are classified into three subpopulations based on distinct morphology: small granulosa cells, intermediate granulosa cells, and a specialized type known as pyriform cells. Pyriform cells are unique to squamates and are conserved across the clade (Goldberg, 1970; Neaves, 1971; Taddei, 1972; Hubert, 1976; Aldridge, 1982; Van Wyk, 1984; Uribe et al., 1996; Callebaut et al., 1997; Ochotorena et al., 2005; Moodley and Van Wyk, 2007; Tumkiratiwong et al., 2012; Lozano et al., 2014; Santos et al., 2015; Afsharzadeh et al., 2015; Hojati et al., 2016; Thongboon et al., 2020, 2022; Xavier et al., 2022; Cruz-Cano et al., 2023; Weberling et al., 2025). Pyriform cells are large compared to the small and intermediate granulosa cells and characterized by a “teardrop” shape, with the narrower end oriented toward the oocyte. They are most prominent during the previtellogenic stages of follicle maturation. Ultrastructural studies suggest that pyriform cells serve as nurse cells because they contain ribosomes, liposomes, mitochondria, and vacuoles within intercellular bridges with the oocyte during later stages of follicle development (Andreuccetti, 1992; Andreuccetti et al., 1978; Bou-Resli, 1974; Filosa and Taddei, 1976; Motta et al., 1995; Neaves, 1971; Taddei and Andreuccetti, 1990). The molecular characteristics of pyriform cells remain poorly defined.

Understanding the molecular organization of the ovary requires tools to visualize gene expression with high spatial resolution. Hybridization chain reaction RNA-fluorescent in situ hybridization (HCR RNA-FISH) is a robust and sensitive method for detecting RNA transcripts by fluorescent microscopy (Choi et al., 2018, 2014, 2010). HCR RNA-FISH has become an increasingly popular method for developmental biology studies (Le et al., 2025; Morrison et al., 2025; Muntzar et al., 2025; Nadolski et al., 2025; Ozekin et al., 2025). In these experiments, the use of a probe set that hybridizes with polyadenylated RNA (poly(A)), a post-transcriptional hallmark of RNA maturation, is commonly used as a positive control for RNA integrity and staining quality (Choi et al., 2016). During HCR RNA-FISH studies of the brown anole ovary, we observed an unexpectedly strong and highly specific poly(A) signal in pyriform cells. Here, we describe the application of poly(A) HCR RNA-FISH to whole-mount ovaries and paraffin sections to assess its utility as a molecular tool for identifying cells with high transcriptional activity or storing abundant poly(A) transcripts.

## Materials and Methods

### Lizards

Brown anole lizards (*Anolis sagrei*) were purchased from Strictly Reptiles, Florida. Anoles were anesthetized by intracelomic injection of 1% buffered tricaine methanesulfonate and then euthanized with an intracelomic injection of 50% tricaine methanesulfonate (Conroy et al., 2009). Animal procedures were approved by the Institutional Animal Care and Use Committee of the University of Texas MD Anderson Cancer Center. Studies were performed consistent with the National Institutes of Health Guide for the Care and Use of Laboratory Animals.

### Ovary Dissection

Ovaries were dissected into phosphate-buffered saline (PBS), removing associated membranes containing pigmented cells. Brightfield images were acquired using a Leica MZ10F dissecting microscope with a Jenoptik Gryphax camera.

### Hybridization chain reaction fluorescent in situ hybridization (HCR RNA-FISH)

#### Wholemount

Ovaries were fixed overnight in 4% paraformaldehyde (PFA) in PBS at 4°C, then washed three times in PBS for 5 minutes each. Tissues were dehydrated through a methanol series (25%, 50%, 75% in PBS and finally 100%, 5 minutes each on ice) and stored in 100% methanol at −20°C. This method was adapted from the Molecular Instruments HCR RNA-FISH (v3.0) protocol for whole-mount mouse embryos (Choi et al., 2016).

Samples were rehydrated through a reverse methanol series (75%, 50%, 25%, PBS, 5 minutes each on ice), then treated with 10 μg/mL Proteinase K for 15 minutes at room temperature. Tissues were then post-fixed in 4% PFA in PBS for 20 minutes at room temperature and washed twice in PBS for 5 minutes each. Samples were then pre-hybridized in Probe Hybridization Buffer (Molecular Instruments) for 30 minutes at 37°C on a nutator. Probes designed and synthesized by Molecular Instruments (v3.0) targeting polyadenylated (poly(A)) RNA were then diluted to a final concentration of 4 nM in Probe Hybridization Buffer, applied to the tissue, then incubated overnight at 37°C.

Following hybridization, samples were washed four times in Probe Wash Buffer (Molecular Instruments) at 37°C for 15 minutes each, then washed twice in 5X saline-sodium citrate with 0.1% Tween-20 (5X SSCT) at room temperature for 5 minutes each. Hairpins labeled with B4 initiators were snap-cooled by heating to 95°C for 90 seconds, then cooled to room temperature in the dark for 30 minutes. Amplification was carried out by incubating tissues in Amplification Buffer (Molecular Instruments) containing 30 nM of each hairpin (60 nM total) overnight at room temperature in the dark on a nutator.

After amplification, samples were washed twice in 5X SSCT for 5 minutes each at room temperature. Samples were then counterstained with 4’,6-diamidino-2-phenylindole (DAPI, 1 μg/mL in 5X SSCT) for 30 minutes at room temperature in the dark. Following staining, samples were washed twice in 5X SSCT for 30 minutes each, then washed a final time in 5X SSCT for 5 minutes at room temperature. Samples were mounted in 5X SSCT with #1.5 coverslips and stored at 4°C in the dark until imaging.

#### Paraffin Sections

Ovaries were fixed overnight in 4% PFA in PBS at 4°C, then washed in PBS and transferred to 70% ethanol. Samples were processed using a Leica TP1020 tissue processor, which carried the tissue through a graded ethanol dehydration series, followed by clearing in Histo-Clear (National Diagnostics HS2001GLL) and infiltration with paraffin. Each of the 12 processing steps was set to 1 hour. Tissues were embedded in paraffin and sectioned at a thickness of 5 µm.

This protocol was adapted from the Molecular Instruments HCR RNA-FISH (v3.0) protocol for Formalin-Fixed Paraffin-Embedded (FFPE) tissue sections (Choi et al., 2016). Slides were heated at 60°C overnight, then deparaffinized in Histo-Clear via three 5-minute washes at room temperature and rehydrated through a graded ethanol series (100% ethanol, 2 X 3 minutes, 95%, 1 X 3 minutes, 70%,1 X 3 minutes, 50%, 1 x 3 minutes, water 1 X 3 minutes). For antigen retrieval, slides were placed in boiling 1X Tris-EDTA buffer (pH 9.0) for 15 minutes on a 95 °C hot plate, then gradually cooled by adding room temperature water every 5 minutes for 20 minutes to bring the solution to ∼45°C. Slides were washed in water for 10 minutes at room temperature, then in PBST (PBS + 0.1% Tween-20) twice for 2 minutes each at room temperature. Slides were pre-hybridized with Probe Hybridization Buffer (Molecular Instruments) at 37°C for 10 minutes in a humidified chamber. Probes designed and synthesized by Molecular Instruments (v3.0) targeting polyadenylated (poly(A)) RNA were then diluted to 1 pmol of probe in 100 μL of Probe Hybridization Buffer per slide, applied to each slide, and incubated overnight at 37°C in a dark, humidified chamber.

Slides were washed at 37°C in a five-step gradient (100% Probe Wash Buffer, 75% probe wash buffer/25% 5X SSCT, 50%/50%, 25%/75%, 100% 5X SSCT) for 15 minutes each step, followed by a final wash in 5X SSCT at room temperature for 5 minutes.

Hairpins labeled with B4 initiators were snap-cooled by heating to 95°C for 90 seconds, then cooled to room temperature in the dark for 30 minutes. For each slide, 15 pmol of each hairpin was added to 100 μL Amplification Buffer (Molecular Instruments), applied to the tissue, and incubated overnight at room temperature in a dark, humidified chamber.

Slides were washed in 5X SSCT for 5 minutes, then counterstained with DAPI (1 μg/mL in 5X SSCT) for 15 minutes at room temperature. Slides were then washed twice in 5X SSCT for 15 minutes at room temperature, washed for a final time for five minutes in 5X SSCT for 5 minutes, and mounted with #1.5 coverslips using 80% glycerol/20% PBS to facilitate later removal of the coverslips for additional staining. Samples were stored at 4°C in the dark until imaging.

#### Fluorescent Microscopy and Image Analysis

Fluorescent images were acquired using an A1 Nikon inverted confocal microscope with a LU-NV laser unit and a DUG GaAsP detector unit. Tiff stacks of the acquired data were processed using Fiji (Schindelin et al., 2012).

Imaris Version 10.2 software (Bitplane, Oxford Instruments) was used to select a ∼17,000 mm^3^ Region of Interest (ROI) on three different wholemount ovaries hybridized with the poly(A) probe for HCR RNA-FISH. These ROIs had a uniform distribution of fluorescent cells and were mostly devoid of blood vessels. Fluorescent cells were identified as a “surfaces” object based on thresholding parameters. The sphericity of each cell was then determined.

#### Histology

Slides previously stained by RNA FISH-HCR were submerged in PBS for 10 minutes to gently float off coverslips without damaging sections. Sections were then stained with hematoxylin and eosin (H&E) following standard procedures and mounted with #1.5 coverslips using Permount (Fisher Scientific, Cat. # SP15-100). H&E-stained sections were imaged using a Nikon Eclipse 80i compound microscope.

## Results

### Organization of follicles in the adult brown anole ovary

Brown anole ovaries showed an organized size gradient of follicles, with the smallest, least-developed white pre-vitellogenic follicles found in a staggered array of increasing size along the anterior surface of the largest, most developed follicle (**Fig. 1A, B**). The most mature follicle was positioned nearest to the reproductive tract organs (**Fig. 1A**). This spatial arrangement reflects the sequential maturation of follicles on each ovary, in which the largest follicle is positioned to be ovulated first (**Fig. 1A**). The second largest follicle is located on the other ovary and should be ovulated next. In the brown anole, there is a left-right alternation of ovulation from the paired ovaries, allowing continuous production of single-egg clutches throughout the breeding season (Crews, 1977; 1980). Upon ovulation, the largest follicle in the contralateral ovary advances toward maturity.

**Fig. 1.**
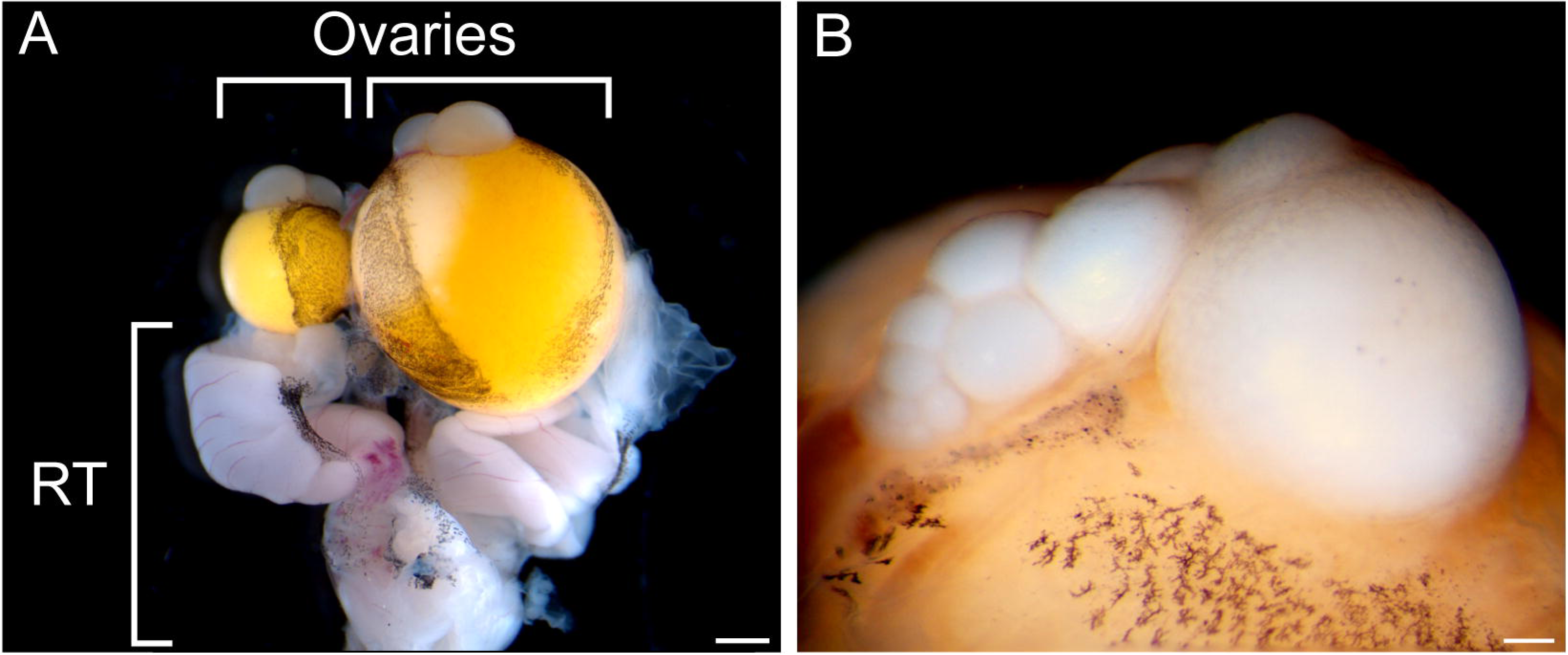
Brown anole ovaries and follicle organization. **A,** Ventral view of breeding season ovaries with intact reproductive tract (RT) organs. Scale bar = 1 mm. **B,** Isolated ovary dissected away from the reproductive tract to show follicle organization. Follicles are organized in a size gradient, with the largest, yolk-filled vitellogenic follicle positioned closest to the RT organs, and progressively smaller, white previtellogenic follicles extending along the anterior end of the ovary. Scale bar = 200 μm.

### Whole-mount poly(A) labeling by HCR RNA-FISH marks a specific somatic cell type in ovarian follicles

We performed HCR RNA-FISH to whole-mount preparations of the adult brown anole ovary to assess RNA integrity and staining quality. We used a probe set targeting polyadenylated RNA (poly(A)), which labels the total pool of mature mRNAs in each cell. This approach provided an opportunity to validate staining performance in brown anole tissues. The poly(A) probe set resulted in robust fluorescent signals (**Fig. 2**). In the absence of the poly(A) probe set there was no signal (**Supp. Fig. 1**).

**Fig. 2.**
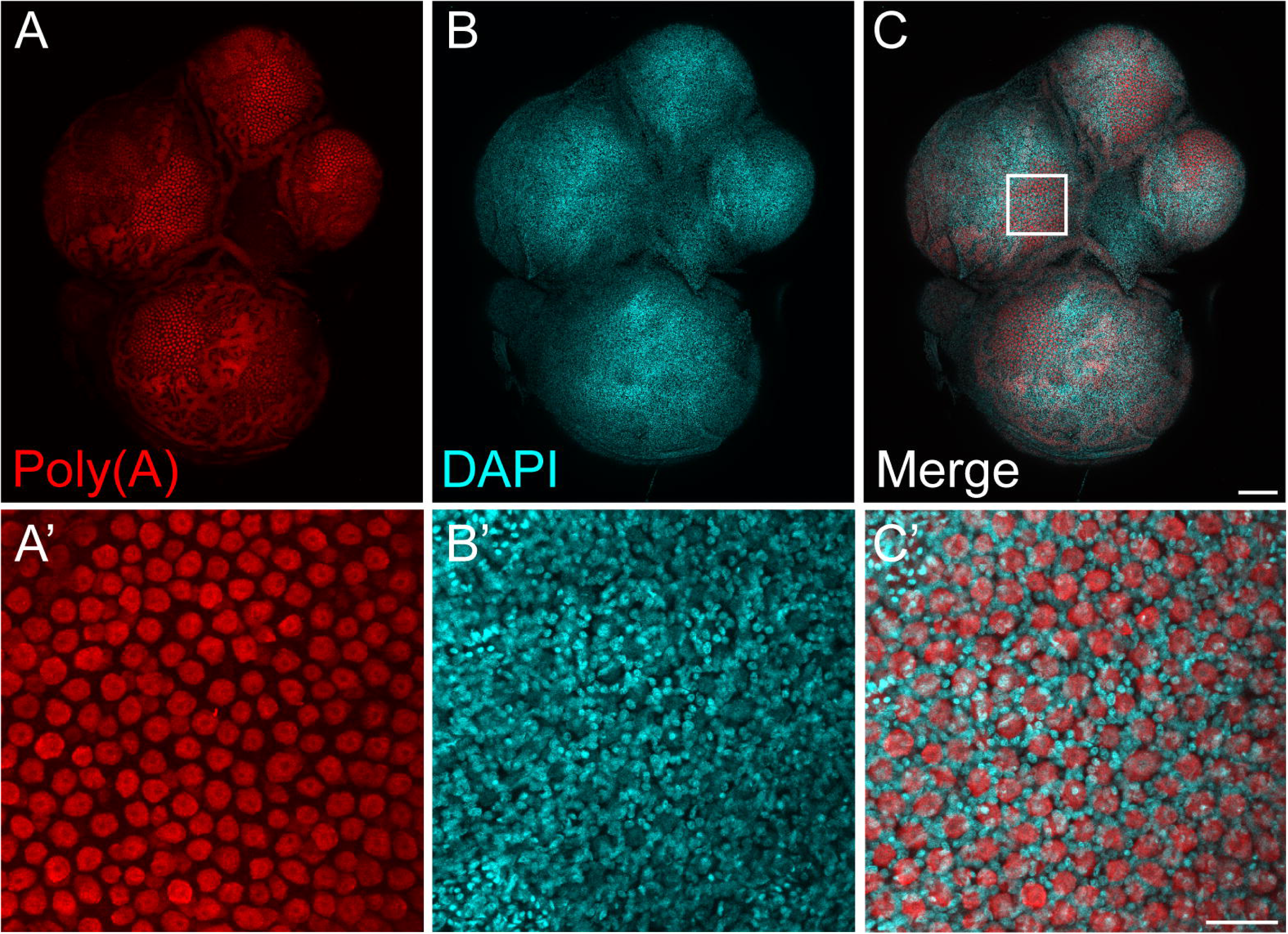
Whole-mount brown anole ovary consisting of previtellogenic follicles stained using HCR RNA-FISH. **A,** Poly(A) (red), **B,** DAPI (cyan), and **C,** merge of A and B. Scale bar = 200 µm. A’-C’) Magnified views of the boxed region outlined in panel C. Scale bar = 10 μm. n=3.

In squamate reptiles, the ovary contains somatic cells that include small and intermediate granulosa cells as well as pyriform cells, which are large, morphologically distinct, teardrop-shaped nurse cells found in previtellogenic follicles. Using the poly(A) probe set, HCR RNA-FISH yielded strikingly bright signals in large cells evenly distributed on the surface of ovarian follicles (**Fig. 2A, A’**). The large size of the cells exhibiting the intense fluorescent signals indicated that they might be pyriform cells. The smaller granulosa cells were present between the putative pyriform cells and exhibited negligible HCR RNA-FISH signal (**Fig. 2C’**). In whole-mount ovaries containing only previtellogenic follicles, poly(A) signal was consistently intense in the presumptive pyriform cells across all previtellogenic follicles, regardless of size (**Fig. 2A)**.

We computationally segmented the highly fluorescent cells from the volumetric dataset obtained from the whole-mount preparations using the surfaces module in Imaris (**Fig. 3A, A’, B, B’, Supplemental Video 1**). A basal view of the fluorescent cells showed that they were not spherical (**Fig. 3C, Sup. Vid. 1**). The side of the cell closest to the follicle had a tapered morphology (**Fig. 3C, Sup. Vid. 1**) and segmentation of individual cells confirmed that they had a teardrop shape (**Fig. 3D, Sup. Vid. 1**). We calculated the sphericity of the fluorescent cells in a region of interest and colored-coded the range of sphericity (**Fig. 3B, B’, Sup. Vid. 1**). A spherical cell will have a sphericity index of 1.0. We confirmed that the fluorescent cells were not spherical (**Fig. 3E, Sup. Vid. 1**). The size and morphology of the poly(A) positive cells confirms that they are pyriform cells.

**Fig. 3.**
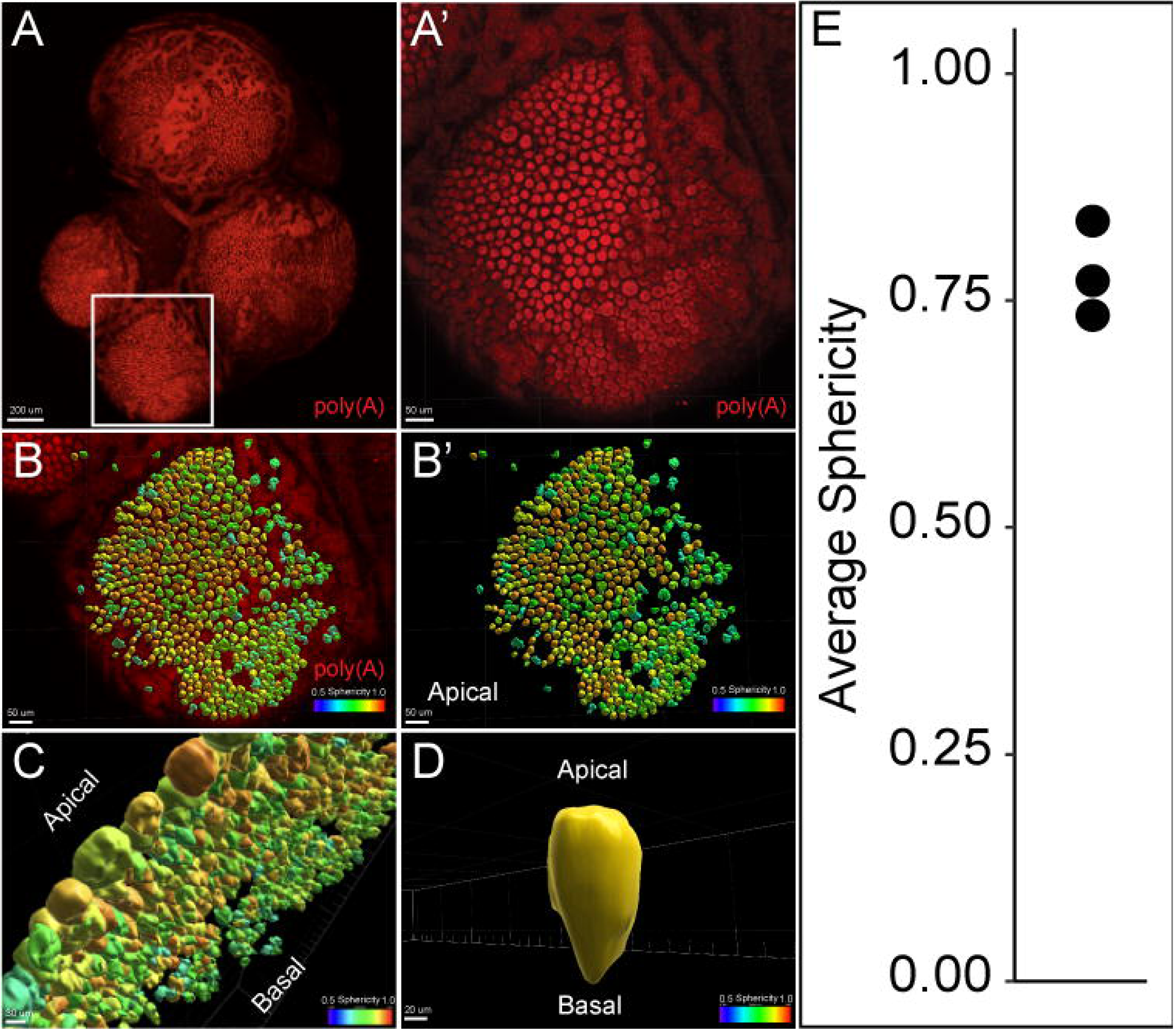
Segmentation of poly(A) HCR RNA-FISH signal in the brown anole ovary. **A,** Surface view of wholemount ovary hybridized with poly(A) probe. **A’,** Higher magnification of boxed area shown in A. **B,** Imaris surface rendering of poly(A)-positive cells. Different colors represent range of cell sphericity. **A’’,** Same as B without fluorescent image.. **B,** Interior view of and **C,** Higher magnification of Imaris surface rendering of poly(A)-positive cells shown from basal (toward the oocyte) view. **D,** Surface rendering of a single poly(A)-positive cell.

DAPI counterstaining revealed weak, diffuse staining in the relatively large nuclei of the pyriform cells (**Fig. 2C’**), whereas the DAPI signal in the smaller nuclei of the adjacent granulosa cells was relatively very strong. These nuclear staining patterns suggest differences in chromatin organization in pyriform cells relative to the surrounding granulosa cell populations. Blood vessels, seen as fluorescent structures in both experimental and no-probe controls due to the strong autofluorescence of lizard erythrocytes (Alibardi, 2021; Storks et al., 2023), were visible in both conditions, confirming red blood cell signal is independent of probe binding (**Sup. Fig. 1A, A’**).

### Paraffin section HCR RNA-FISH using poly(A) probe set selectively marks pyriform cells in the brown anole ovary

HCR RNA-FISH of paraffin sections of the ovary using the poly(A) probe set further confirmed that the intensely staining cells were pyriform cells in the previtellogenic follicles, whereas the small and intermediate granulosa cells had minimal signal (**Fig. 4**). The intense fluorescent signal in pyriform cells highlighted their unique inverted teardrop-shaped morphology (**Fig. 4A’, C’**). Previtellogenic follicles showed robust poly(A) signal in pyriform cells (**Fig. 4A, A’, C, C’)**, whereas late vitellogenic follicles that lack pyriform cells exhibited minimal signal (**Fig. 4E, E’, G, G’**). This was consistent across biological replicates (n = 7), supporting the utility of poly(A) as a molecular marker of pyriform cells. As found in the wholemount studies, pyriform nuclei had weak DAPI-stained nuclei, compared to strong DAPI staining in small and intermediate granulosa cell nuclei (**Fig. 4B’, C’**).

**Fig. 4.**
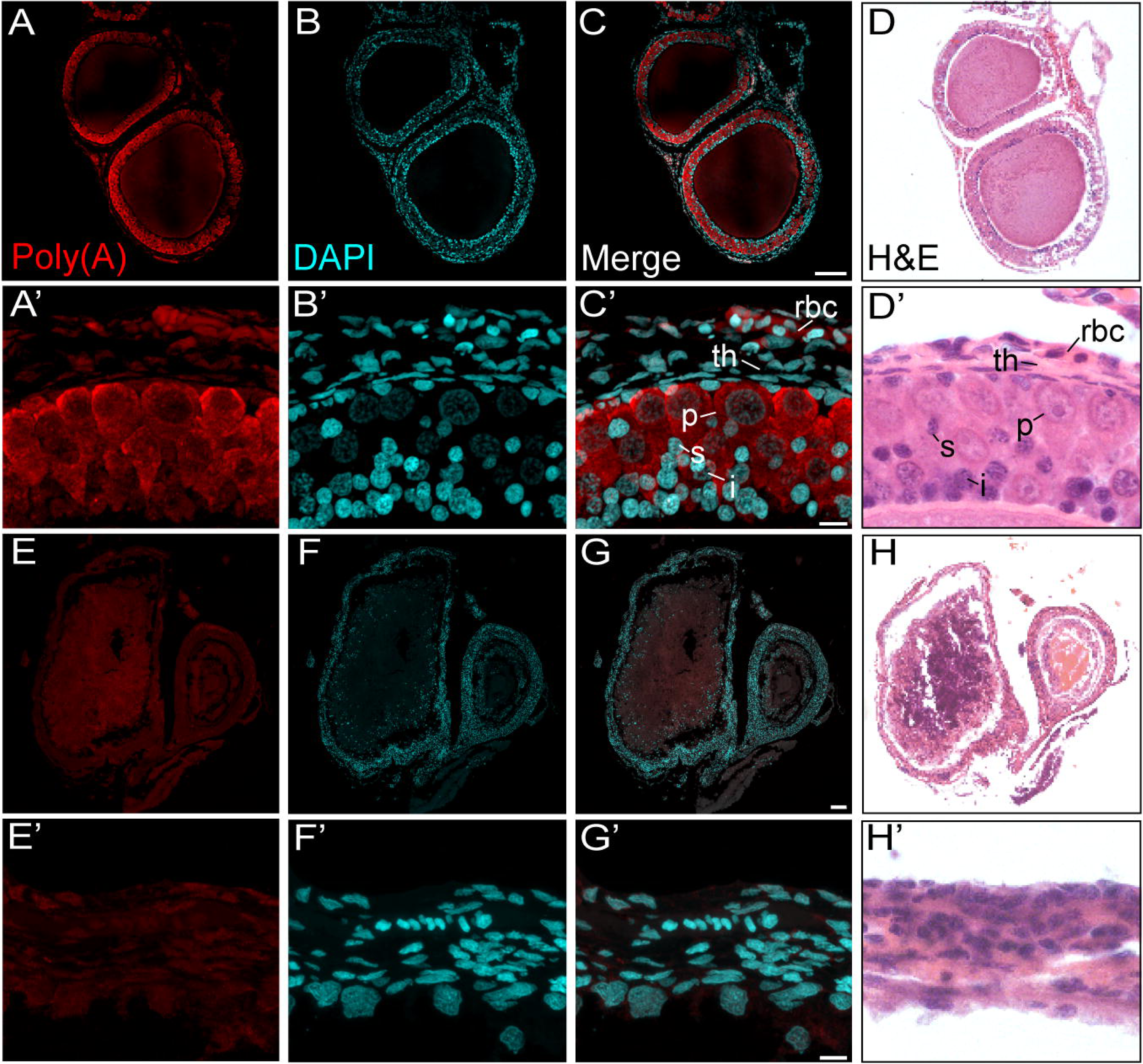
HCR RNA-FISH and H&E of paraffin sections of the brown anole ovary. **A-C,** Cross section of ovary with previtellogenic follicles. **A,** Poly(A) probe (red), **B,** DAPI (cyan), **C,** Merge of A and B. Scale bar = 100 µm. **D,** Same section shown in A-C, stained for H&E. **A’-C’,** Higher magnification of A-C showing somatic cell layers adjacent to oocyte. **A’,** Poly(A) probe (red), **B’,** DAPI (cyan), **C’,** Merge of A and B. i, intermediate granulosa cell, p, pyriform cell, rbc, red blood cell, s, small granulosa cell, th, theca cell. Scale bar = 10 µm. **D’,** Comparable region as A’-C’, stained for H&E. **E-G,** Cross section of ovary with vitellogenic follicles. Scale bar = 100 µm. **H,** Same section shown in E-G, stained for H&E. **E’–G’,** Higher magnification of E-G showing somatic cell layers adjacent to oocyte. **E’,** Poly(A) probe (red), **F’,** DAPI (cyan), **G’,** Merge of A and B. Scale bar = 10 µm. **H’,** Comparable region as E’-G’, stained for H&E. n=8.

To facilitate interpretation of the HCR RNA-FISH results, the same paraffin sections analyzed by HCR RNA-FISH were subsequently stained with H&E, providing detailed morphological context for identifying granulosa cell types and follicle organization. H&E staining revealed the somatic cell organization of developing follicles, including the granulosa cell population, which in reptiles consists of three morphologically distinct types in previtellogenic follicles: small granulosa cells, intermediate granulosa cells, and pyriform cells (**Fig. 4D, D’**). Pyriform cells are most prominent at previtellogenic stages, appearing as large, teardrop-shaped cells with weakly-staining nuclei. In late-stage follicles, pyriform cells are no longer present, as they have undergone apoptosis by this stage (Santos et al., 2020) (**Fig. 4H, H’)**. For both previtellogenic and vitellogenic stage follicles, the outer boundary of each follicle consisted of at least one layer composed of elongated, flattened thecal cells (**Fig. 4D’, H’)**. The ovary is vascularized, thus in sections, nucleated red blood cells, which are typical of reptiles, are scattered throughout the thecal layer (**Fig. 4D’)**. These red blood cells exhibit a distinct, round, flattened morphology and eosinophilic cytoplasmic staining that differentiates them from theca cells. This histological reference provides a clear morphological framework for identifying cell types and organization in corresponding poly(A) HCR RNA-FISH sections.

## Discussion

In situ hybridization is a fundamental tool used by developmental biologists to visualize the location of specific nucleic acid sequences in cells and tissues (Bauman et al., 1980; Gall and Pardue, 1969; Tautz and Pfeifle, 1989). HCR RNA-FISH is gaining in popularity as a method for assessing spatial patterns of gene expression in whole-mount and sectioned specimens (Le et al., 2025; Morrison et al., 2025; Muntzar et al., 2025; Nadolski et al., 2025; Ozekin et al., 2025). While establishing HCR RNA-FISH for our studies of the brown anole, we used the poly(A) probe set to assess the method in whole-mount and paraffin-sectioned ovaries. To our surprise, we found very strong fluorescent signals specifically in pyriform cells of previtellogenic follicles, while other cell types had negligible signal. Our findings suggest that a poly(A) probe set can be used as a cell type-specific marker in certain tissues.

The ovary contains germ cells and multiple somatic cell types organized into follicles and other tissues essential for reproduction and physiology. Among these somatic cell populations, nurse cells play an important role in provisioning the developing oocyte. In squamate reptiles, this role is fulfilled by pyriform cells, a large, morphologically distinct granulosa cell type unique to the clade and highly conserved across species (Goldberg, 1970; Neaves, 1971; Taddei, 1972; Hubert, 1976; Aldridge, 1982; Van Wyk, 1984; Uribe et al., 1996; Callebaut et al., 1997; Ochotorena et al., 2005; Moodley and Van Wyk, 2007; Tumkiratiwong et al., 2012; Lozano et al., 2014; Santos et al., 2015; Afsharzadeh et al., 2015; Hojati et al., 2016; Thongboon et al., 2020, 2022; Xavier et al., 2022; Cruz-Cano et al., 2023; Weberling et al., 2025). Prior to the current study, there were no molecular markers of squamate ovarian pyriform cells.

Although pyriform cells are functionally analogous to the well-studied ovarian nurse cells of *Drosophila*, there are important differences in their morphology and connectivity to the oocyte. In *Drosophila*, nurse cells are joined to the oocyte via numerous wide protrusions, facilitating rapid “dumping” of nutrients and cytoplasmic contents into the oocyte during late oogenesis (Riparbelli et al., 2022). In contrast, squamate pyriform cells connect to the oocyte through individual thin intercellular bridges (Andreuccetti, 1992; Andreuccetti et al., 1978; Bou-Resli, 1974; Filosa and Taddei, 1976; Motta et al., 1995; Neaves, 1971; Taddei and Andreuccetti, 1990), which may limit the rate of transfer and suggest a more gradual or selective delivery of materials. This structural difference may underlie distinct timing or regulatory mechanisms of resource delivery in squamate follicles compared to insect systems.

The intense fluorescent signal found throughout the cytoplasm of the pyriform cells highlighted their unique teardrop morphology. The tip of the pyriform cell is the location where the intercellular bridge forms with the developing oocyte. We found that pyriform cells were distributed homogeneously on the surface of each follicle. This suggests that the corresponding intercellular bridges with the maturing oocyte are also homogeneously distributed, perhaps facilitating the equal distribution of nurse cell products into the oocyte. In the green anole (*Anolis carolinensis*), a close relative of *Anolis sagrei*, these intercellular bridges are relatively thin, such that the connection with the oocyte in histology only appears as a slight “puckering” of the membrane between the oocyte and the somatic cell layers (Neaves, 1971). This is in contrast to other squamates, such as *Tropidurus* lizards, which have a relatively large and visible hole in the oolemma (Silva et al., 2018). This indicates that, although pyriform cell morphology and function are conserved across squamates, there is some variation in the morphology of intercellular bridges.

The strong fluorescent signal found in pyriform cells using the poly(A) probe suggests that these cells contain relatively large amounts of polyadenylated RNA. This suggests that pyriform cells have exceptionally high transcriptional output, reservoirs for mature mRNAs, or both. This is consistent with ultrastructural evidence showing that pyriform cells are rich in ribosomes, mitochondria, and other biosynthetic organelles, especially in the intercellular bridge linking them to the oocyte (Motta et al., 1995).

Cell type-specific expression has been reported using poly(A) probes in other in situ hybridization methods (Markovic et al., 1992; Pringle et al., 1989), suggesting that our findings are not unique to squamate ovaries. It is possible that other cell types with high transcriptional activity or large stores of mature mRNA, such as nurse cells in other taxa, specialized secretory cells, embryonic stem cells, or cancer cells, may exhibit similar staining patterns (Capco and Jeffery, 1979; Kobayashi et al., 1988; Efroni et al., 2008; Kotsantis et al., 2016; Percharde et al., 2017; Bradner et al., 2017; Connors et al., 2024).

Despite their important function and conservation across squamate species, pyriform cells have not been well characterized, and no molecular marker that remains consistent across stages of follicle development has been defined. Here, we identify a reliable molecular marker for pyriform cells, enabling visualization of their distribution in both whole-mount and sectioned tissue. Beyond its immediate application to reptile biology, this approach opens opportunities to investigate nurse cell transcriptional specialization, intercellular transport, and follicular organization in a broad range of species, thereby contributing to comparative and evolutionary analyses of ovarian structure and function.

## Acknowledgements

We thank the members of our lab for helpful comments. This research was supported by National Institutes of Health (NIH) grant HD113569, Sigma Xi Grant in Aid of Research G20230315-4769, and the Ben F. Love Endowment to R.R.B. Confocal microscopy was supported by NIH shared instrumentation grant OD024976. Veterinary resources were supported by NIH Grant CA16672.

**Supplemental 1.** Whole-mount brown anole ovary HCR RNA-FISH negative control. **A,** No probe (red), **B,** DAPI (cyan), and **C,** Merge of A and B. Scale bar = 200 µm. **A’-C’**, Magnified views of the boxed region in panel C. p, pyriform cell, rbc, red blood cell. Scale bar = 10 μm. n = 3.

**Supplemental Video 1.** Video showing 3D morphology of pyriform cells through sequential layering and different views of whole-mount HCR poly(A) staining and surface rendering of pyriform cells. The first rotation shows only the whole-mount HCR. Next, the pyriform surface rendering is superimposed onto the HCR stain. The zoom-in shows a basal view of the pyriform cells, highlighting their teardrop morphology. Finally, we show a single pyriform cell shape in 3D.

